## Supporting Information for "Children exhibit a developmental advantage in the offline processing of a learned motor sequence"

#### Appendix 1: Participant characteristics, sleep and vigilance (Experiments 1 and 2)

Group means for the measures of both Experiments 1 and 2 are provided in Tables 1 and 2 in the main text and results from the corresponding statistical analyses can be found in Tables S1 and S2 below. In brief, significant age group differences were observed for gender distribution, morningness/eveningness preference, sleep duration of the night prior to participation and subjective levels of sleepiness at the time of testing for both experiments. Additionally, the time of testing and objective levels of alertness were different among age groups in Experiment 1 and 2, respectively. These age group differences can largely be considered reflective of the convenience sample in the current research as well as known lifespan differences in a subset of these measures. We elaborate on these differences in the subsequent paragraphs.

The adult groups showed a gender distribution skewed towards more female than male participants. This group difference can largely be linked to our convenience sample. The potential impact of these differences in gender distribution across our age groups on our primary measures (i.e., learning magnitude, micro-online gains, and micro- and macro-offline gains) was assessed. Results from the corresponding gender by age group interactions revealed no significant effects (all $p > 0.08$).

Older adults indicated a greater preference for mornings than the other age groups. This is in line with previous studies indicating that circadian preferences shift towards the morning with increasing age in adulthood[1,2].

In Experiment 1, participants were instructed to complete the experiment between 9 am and 7 pm to avoid testing early in the morning or late at night when performance is more likely to be impacted by circadian influences. Nonetheless, young adults completed the experimental session later in the day as compared to the other age groups, potentially due to work and/or family responsibilities during the day. In Experiment 2, participants were given specific instructions regarding the timing of the sessions, decreasing the flexibility in the time windows for task completion and thus there were no differences among age groups.

Self-reported sleep duration was longer in children in both experiments and in adolescents in Experiment 2 as compared to the adult groups. Furthermore, older adults reported a shorter sleep duration in comparison to the other age groups in Experiment 1. These findings are consistent with a known decrease in total sleep time with age[3,4]. There were no differences between the two nights in Experiment 2.

Sleep quality during the night prior to the experimental sessions was comparable among age groups and experimental nights in both experiments. Based on previous literature, one would expect older adults to report decreased sleep quality[5]. However, as Vitiello et al.[6] found a mismatch between self-reported and objectively measured sleep quality, it is possible that our older adults overestimated the quality of their sleep.

Lastly, and unexpectedly, older adults self-reported a higher alertness than the other age groups in both experiments, with a group by session interaction in Experiment 2. We speculate that, and analogous to sleep quality, older adults underestimated their levels of sleepiness at the time of testing. This is supported by the fact that the older adults were not different from the adolescents and young adults in their performance on the PVT – an objective measure of vigilance - in Experiment 2. Children were slower on the PVT as compared to the other age groups, which is in line with previous research[7].

***Table S1:*** *Participant characteristics in Experiment 1.*

| **Variable** | **df** | **$F/X^2$** | **p** | **$\eta^2/\phi c$** |
|---|---|---|---|---|
| Gender *(n)* | 3, 130 | 13.27 | 0.004* | 0.319 |
| M/E Preference | 3,129 | 5.98 | < 0.001 * | 0.125 |
| Time of testing (SD in min) | 3, 129 | 6.46 | < 0.001* | 0.133 |
| Sleep quality | 3, 129 | 1.98 | 0.120 | 0.045 |
| Sleep duration (min) | 3, 129 | 19.49 | < 0.001* | 0.317 |
| SSS score | 3, 129 | 3.05 | 0.031* | 0.068 |

*Results from statistical analyses assessing group differences in gender, Morningness/Eveningness (M/E) preference, time of testing, sleep quality and quantity, and a subjective vigilance measure. Gender reports chi square statistics and the Cramer's V effect size, whereas all other variables list F-values and eta-squared effect sizes. Significant values are marked with an asterisk. Df = degrees of freedom; $\eta^2$ = eta squared; SSS = Stanford Sleepiness Scale[8]. Corresponding group means are provided in Table 1 of the main text and depicted in Figure S1 as part of this Supporting Information document. Figure S1 also indicates significant pairwise comparisons that were conducted as follow-ups to significant contrasts shown above.*

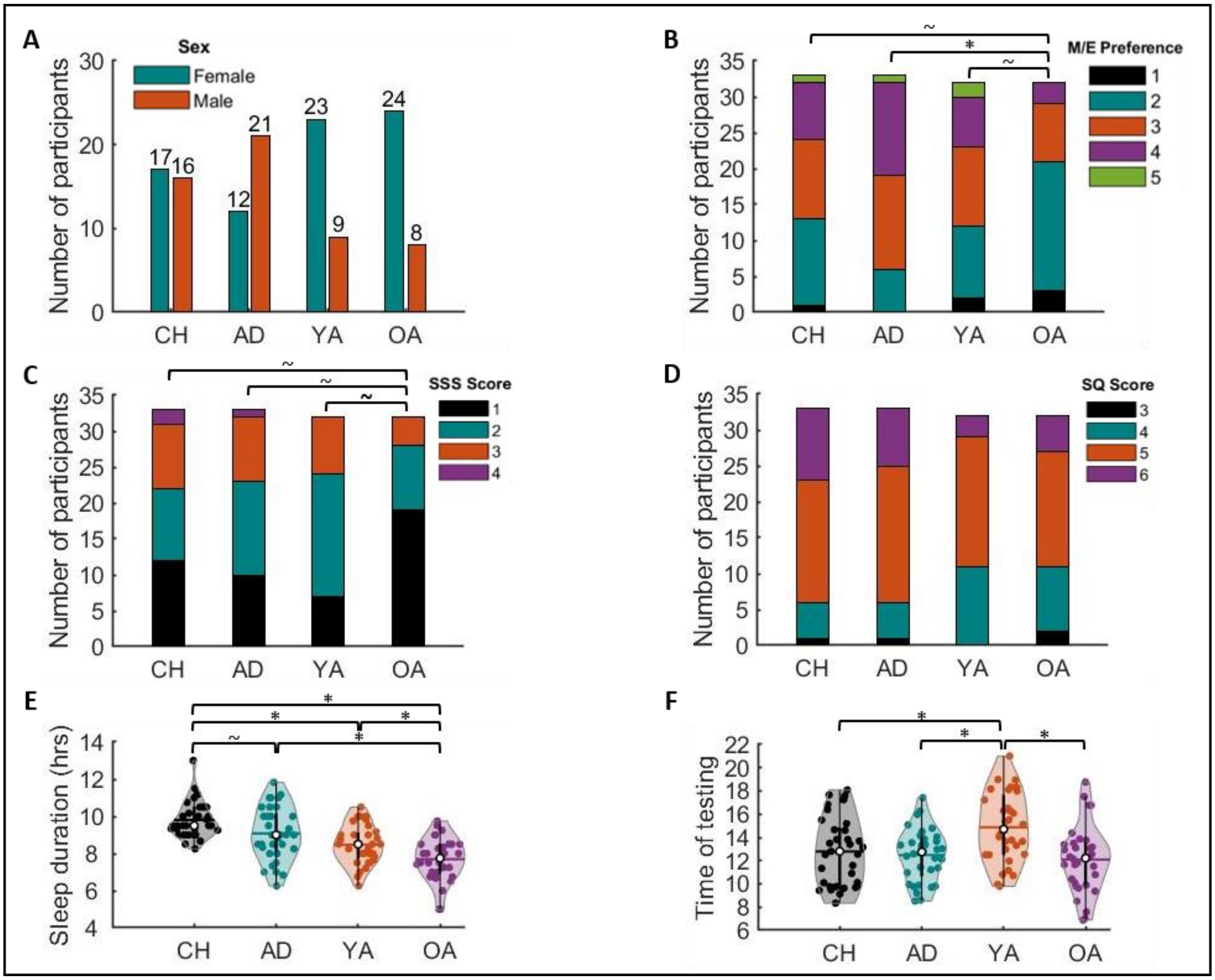

**Figure S1**: *Experiment 1.* ***A****) Gender distribution within the four age groups.* ***B****) Morningness/Eveningness (M/E) preference (1 = extreme morning, 2 = moderate morning, 3 = intermediate, 4 = moderate evening, 5 = extreme evening);* ***C)*** *Subjective vigilance at time of testing (Stanford Sleepiness Scale score*[8]*);* ***D****) Sleep quality of the night prior to testing, scored on a 6-point Likert scale (1 = very bad; 2 = bad; 3 = fairly bad, 4 = fairly well, 5 = well, 6 = very well);* ***E****) Sleep duration of night prior to testing;* ***F****) Time of testing specified in 24-hour time notation. Results from corresponding statistical analyses are presented in Table S1 above. For violin plots in Panels E and F, shaded regions represent the kernel density estimate of the data, colored circles depict individual data, open circles represent group medians, and the horizontal lines depict group means*[9]*. * indicates significant differences ($p < 0.05$) and ~ indicates marginally significant differences ($0.05 < p < 0.1$) between pairs of groups.*

***Table S2**: Participant characteristics in Experiment 2.*

| Variable | df | F/X² | p | η²/ ϕc |
|---|---|---|---|---|
| *A. Gender (n)* | 3,107 | 6.72 | 0.081 | 0.249 |
| *B. M/E Preference* | 3,107 | 5.16 | 0.002* | 0.130 |
| *C. Time of testing* | | | | |
| Session | 1.41,146.63 | 814.80 | < 0.001* | 0.887 |
| Group | 3,104 | 0.47 | 0.702 | 0.013 |
| Session x group | 4.23,146.63 | 1.42 | 0.229 | 0.039 |
| *D. Sleep quality* | | | | |
| Night | 1,103 | 2.37 | 0.127 | 0.023 |
| Group | 3,103 | 1.06 | 0.371 | 0.030 |
| Night x group | 3,103 | 1.71 | 0.171 | 0.047 |
| *E. Sleep duration (min)* | | | | |
| Night | 1,103 | 0.01 | 0.913 | 0.00 |
| Group | 3,103 | 31.94 | < 0.001* | 0.482 |
| Night x group | 3,103 | 1.64 | 0.186 | 0.045 |
| *F. SSS score* | | | | |
| Session | 2,206 | 1.75 | 0.176 | 0.017 |
| Group | 3,103 | 3.56 | 0.017* | 0.094 |
| Session x group | 6,206 | 2.84 | 0.011* | 0.076 |
| *G. PVT score* | | | | |
| Session | 1.73,179.40 | 1.05 | 0.343 | 0.010 |
| Group | 3,104 | 4.61 | 0.005* | 0.117 |
| Session x group | 5.18,179.40 | 1.79 | 0.115 | 0.049 |

*Results from statistical analyses of gender, Morning/Eveningness (M/E) preference, time of testing, sleep quality and quantity, and subjective and objective vigilance measures of the four age groups. Gender reports chi square statistics and the Cramer's V effect size, whereas all other variables list F-values and eta-squared effect sizes. Significant values are marked with an asterisk. Df = degrees of freedom; $\eta^2$ = eta squared; SSS = Stanford Sleepiness Scale[8]. PVT = psychomotor vigilance task[10]. Corresponding group means are provided in Table 2 of the main text and depicted in Figure S2 as part of this Supporting Information. Figure S2 also indicates significant pairwise comparisons that were conducted as follow-ups to significant contrasts shown above.*

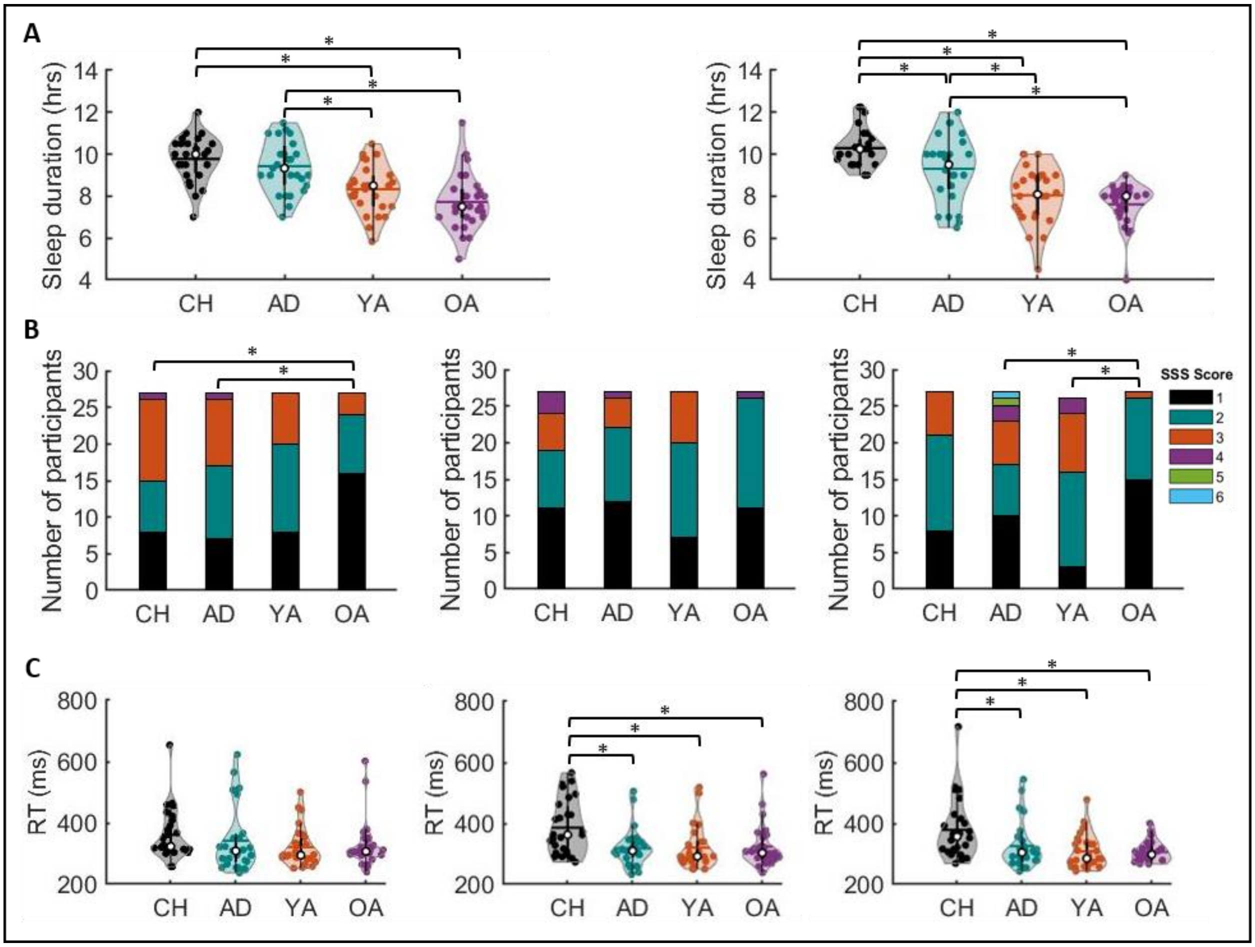

***Figure S2**: Experiment 2.* ***A**) Sleep duration for the night preceding experimental sessions on Day 1 (left) and Day 2 (right).* ***B**) Subjective vigilance[8] at time of testing for session 1 (left), session 2 (middle) and session 3 (right);* ***C)** Objective vigilance, assessed by the Psychomotor Vigilance Task (PVT[10]), at time of testing for session 1 (left), session 2 (middle) and session 3 (right; PVT RT). Results from corresponding statistical analyses are presented in Table S2 above. For all violin plots in Panels A and C, shaded regions represent the kernel density estimate of the data, colored circles depict individual data, open circles represent group medians, and the horizontal lines depict group means[9]. * indicates significant differences between pairs of groups ($p < 0.05$).*

#### Appendix 2: Assessment of learning dynamics with non-normalized performance measures (Experiments 1 and 2)

Results presented in the main text were based on the normalized performance outcomes, adjusted for baseline performance on the pre-learning random run. For completeness, exploratory analyses were also performed on the non-normalized performance outcomes.

Non-normalized performance measures on the SRT task of Experiments 1 and 2 are depicted in Supplemental Figures S3 and S4 and results from the corresponding statistical analyses are provided in Supplemental Tables S3 and S4, respectively. In brief, absolute accuracy remained stable across blocks of practice for all task runs of both experiments, as evidenced by no main effects of block. Furthermore, similar changes across task blocks were found among groups (i.e., no groups x block interaction effects), but the overall performance levels were significantly different among age groups (i.e., group main effect). Specifically, pairwise follow-up comparisons indicated that for certain task runs of the two experiments, children were less accurate than adolescents (i.e., all task runs of Experiment 1, except for the post-learning random), young adults (all runs of Experiment 1 plus pre-learning random and test of Experiment 2), and older adults (all task runs of both experiments except for the post-learning random of Experiment 2).

Absolute response times (RTs) on the pre-learning random revealed a group effect but no block or group x block interaction effect in both experiments. During the training run of both experiments, RTs significantly decreased across practice blocks, as shown by a main effect of block, and the overall RT was significantly different among age groups. Furthermore, in Experiment 1, no group x block interaction effect was found and thus the decrease in RT across the training blocks was similar among age groups. In Experiment 2, however, results revealed a marginally significant group x block interaction, indicating that the change in performance across training blocks tended to be different among groups. During the post-learning test run of both experiments, there was no significant block effect, indicating a performance plateau was reached. This plateau was reached by all groups (i.e., no group x block interaction effect), but the age groups reached significantly different performance levels (i.e., group main effect). Similarly, the post-learning random run showed no block or group x block interaction effects but did reveal significant overall performance differences among groups. Pairwise follow-up comparisons for all the main effects of group indicated that the children were significantly slower than the other age groups for all task runs. Furthermore, the adolescents were slower than the young adults for all task runs of Experiment 2 except for the post-learning test. And lastly, the older adults were slower than the young adults for certain task runs of Experiment 1 (i.e., pre-learning random and post-learning test) and all task runs of Experiment 2.

In summary, and as expected, children were less accurate and slower on the task as compared to the other age groups. Results corresponding to these changes in non-normalized performance metrics across blocks of practice were largely consistent with those in the main text on normalized data.

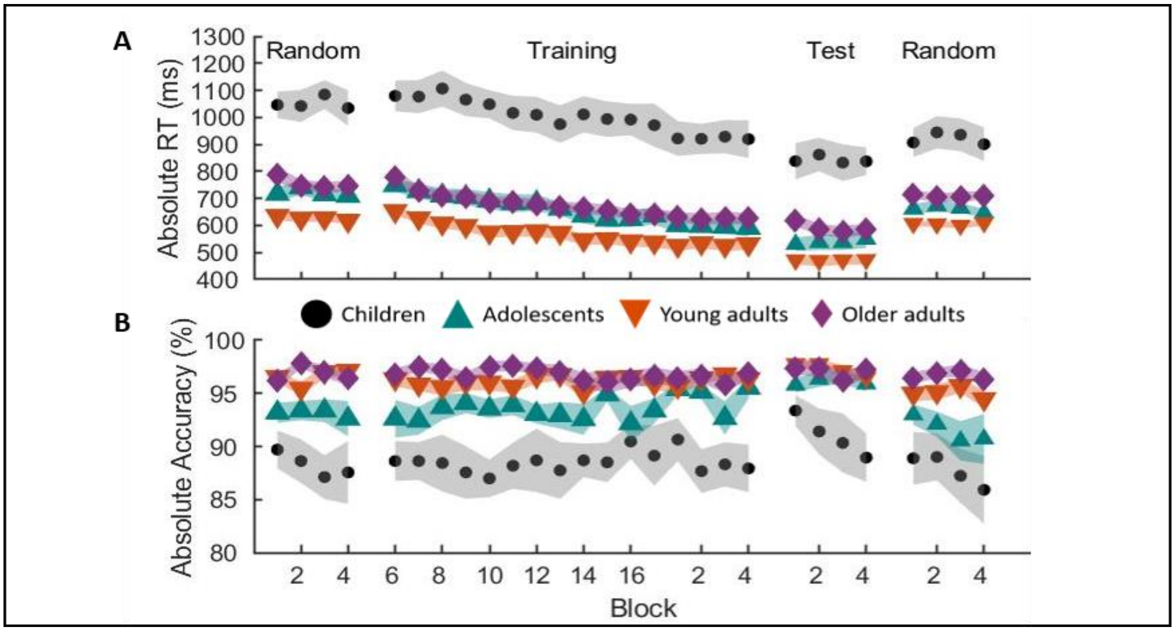

***Figure S3****: Absolute reaction time (Panel A) and absolute accuracy (Panel B) for all task runs in Experiment 1. Shaded regions = SEM. Results from the corresponding statistical analyses are shown in Table S5 below. Behavioral data normalized to the pre-learning pseudo-random run are in Figure 2 in the main text.*

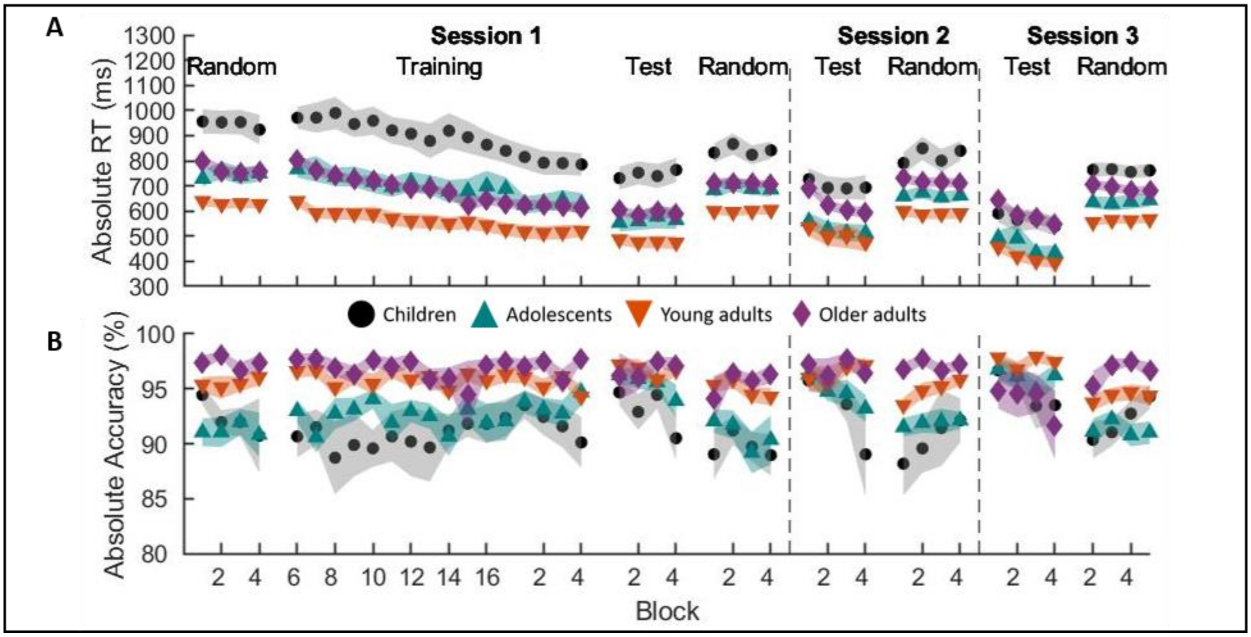

***Figure S4****: **A**) Absolute reaction time in milliseconds for all tasks and sessions. **B**) Absolute accuracy as a percentage of correct keypresses for all tasks and sessions. Shaded regions = SEM. Results from the corresponding statistical analyses are shown in Table S4. Behavioral data normalized to the pre-learning pseudo-random run are in Figure 4 in the main text*

***Table S3**: Absolute (raw) task performance in Experiment 1.*

| | **RT** | | | | **Accuracy** | | | |
|---|---|---|---|---|---|---|---|---|
| **Effect** | **df** | **F** | **p** | **Part η²** | **df** | **F** | **p** | **Part η²** |
| *A. Pre-learning random* | | | | | | | | |
| Block | 2.18,275.25 | 1.42 | 0.242 | 0.011 | 2.01,253.07 | 0.15 | 0.864 | 0.001 |
| Group | 3,126 | 33.12 | <0.001* | 0.441 | 3,126 | 16.23 | <0.001* | 0.279 |
| B x G | 6.55,275.25 | 1.55 | 0.155 | 0.036 | 6.03,253.07 | 0.71 | 0.646 | 0.017 |
| *B. Training* | | | | | | | | |
| Block | 7.21,908.06 | 44.56 | <0.001* | 0.261 | 6.98,879.82 | 0.42 | 0.890 | 0.003 |
| Group | 3,126 | 28.46 | <0.001* | 0.404 | 3,126 | 15.86 | <0.001* | 0.274 |
| B x G | 21.62,908.06 | 1.36 | 0.127 | 0.031 | 20.95,879.82 | 0.75 | 0.778 | 0.018 |
| *C. Test* | | | | | | | | |
| Block | 1.72,216.86 | 0.32 | 0.693 | 0.003 | 2.22,279.79 | 1.81 | 0.160 | 0.014 |
| Group | 3,126 | 21.61 | <0.001* | 0.340 | 3,126 | 10.15 | <0.001* | 0.195 |
| B x G | 5.16,216.87 | 0.63 | 0.679 | 0.015 | 6.66,279.79 | 0.92 | 0.485 | 0.022 |
| *D. Post-learning random* | | | | | | | | |
| Block | 2.49,314.13 | 1.08 | 0.351 | 0.008 | 1.97,248.19 | 1.61 | 0.203 | 0.013 |
| Group | 3,126 | 19.49 | <0.001* | 0.317 | 3,126 | 8.57 | <0.001 | 0.170 |
| B x G | 7.48,314.126 | 1.12 | 0.352 | 0.026 | 5.91,248.19 | 0.59 | 0.737 | 0.014 |

*Results from statistical analyses of raw response time (RT) and accuracy measures for all task runs of Experiment 1. Significant values are marked with an asterisk. Df = degrees of freedom; part $\eta^2$ = partial eta squared; B x G = block x group interaction.*

***Table S4:*** *Absolute (raw) task performance of session 1 in Experiment 2.*

| | RT | | | | Accuracy | | | |
|---|---|---|---|---|---|---|---|---|
| **Effect** | **df** | **F** | **p** | **Part η²** | **df** | **F** | **p** | **Part η²** |
| *A. Pre-learning random* | | | | | | | | |
| Block | 1.69,175.24 | 1.08 | 0.332 | 0.010 | 1.84,191.60 | 0.35 | 0.687 | 0.003 |
| Group | 3,104 | 17.10 | <0.001* | 0.330 | 3,104 | 10.13 | <0.001* | 0.226 |
| B x G | 5.06,175.24 | 1.37 | 0.239 | 0.038 | 5.53,191.60 | 0.69 | 0.645 | 0.020 |
| *B. Training* | | | | | | | | |
| Block | 6.80,706.71 | 34.62 | <0.001* | 0.250 | 6.42,667.37 | 0.86 | 0.532 | 0.008 |
| Group | 3,104 | 15.38 | <0.001* | 0.307 | 3,104 | 7.03 | <0.001* | 0.169 |
| B x G | 20.39,706.71 | 1.56 | 0.055 | 0.043 | 19.25,667.37 | 1.05 | 0.402 | 0.029 |
| *C. Test* | | | | | | | | |
| Block | 2.59,269.64 | 0.28 | 0.808 | 0.003 | 2.23,231.98 | 2.15 | 0.113 | 0.020 |
| Group | 3,104 | 11.26 | <0.001* | 0.245 | 3,104 | 3.62 | 0.016* | 0.095 |
| B x G | 7.78,269.64 | 1.19 | 0.304 | 0.033 | 6.69,231.98 | 1.57 | 0.149 | 0.043 |
| *D. Post-learning random* | | | | | | | | |
| Block | 2.34,243.01 | 1.84 | 0.154 | 0.017 | 2.10,218.70 | 1.83 | 0.161 | 0.017 |
| Group | 3,104 | 15.64 | <0.001* | 0.311 | 3,104 | 7.40 | <0.001* | 0.176 |
| B x G | 7.01,243.01 | 0.90 | 0.509 | 0.025 | 6.31,218.70 | 0.82 | 0.559 | 0.023 |

*Results from statistical analyses of raw response time (RT) and accuracy measures for all task runs of session 1 of Experiment 2. Significant values are marked with an asterisk. Df = degrees of freedom; part $\eta^2$ = partial eta squared; B x G = block x group interaction.*

#### Appendix 3: Assessment of learning magnitude and micro-learning with non-normalized data (Experiment 1)

The dependent measures of learning-magnitude (assessing sequence-specific learning) as well as micro-offline and -online performance changes were computed with *normalized* RT data in the main text. Here, we present results from the same analyses but with these performance indices computed with non-normalized response time (RT) data.

##### *3.1 Learning Magnitude*

The difference in non-normalized RT between the post-learning test (averaged across the 4 test blocks) and the post-learning random (average across the 4 random blocks) was divided by the non-normalized RT in the post-learning random run (averaged across the 4 blocks). A one-way ANOVA revealed a significant group effect ($F_{(3,129)} = 5.440$, $p = 0.002$, $\eta^2 = 0.115$; see Figure S5). Follow-up comparisons revealed that a significantly smaller sequence-specific learning magnitude was observed in children as compared to adolescents ($p = 0.014$, $G = 0.714$) and young adults ($p = 0.001$, $G = 0.970$). This finding is consistent with the results based on the normalized data presented in the main text.

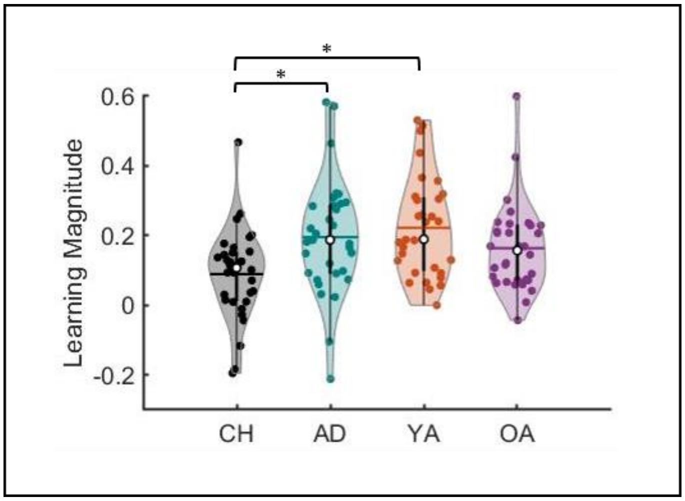

***Figure S5.*** *Average learning magnitude on non-normalized RT per age group. Shaded regions represent the kernel density estimate of the data, colored circles depict individual data, open circles represent group medians, and the horizontal lines depict group means[9]. * $p < 0.05$ for pairwise group comparisons.*

##### *3.2. Micro-learning across sequential training*

Micro-online gains were computed as the difference in averaged non-normalized performance between the correct keypresses of the first and last sequence repetitions within each block. Results of the mixed ANOVA indicated significant block ($F_{(10.002,1240.225)} = 2.312$, $p = 0.011$, partial $\eta^2 = 0.018$) and group ($F_{(3,124)} = 8.280$, $p < 0.001$, partial $\eta^2 = 0.167$) main effects, as well as a significant block x group interaction ($F_{(30.01,1240.23)} = 1.553$, $p = 0.029$, partial $\eta^2 = 0.036$; Figure S6A). Post-hoc pairwise comparisons on group differences indicated significantly smaller (i.e., more negative) micro-online gains in children as compared to adolescents ($F_{(1,62)} = 7.204$, $p = 0.009$, partial $\eta^2 = 0.104$; Figure S6C), young ($F_{(1,61)} =13.610$, $p < 0.001$, partial $\eta^2 = 0.182$) and older ($F_{(1,61)} = 11.952$, $p = 0.001$, partial $\eta^2 = 0.164$) adults.

Micro-offline gains were calculated as the difference in the averaged non-normalized RT between the correct keypresses of the last sequence repetition of one block and the first sequence repetition of the subsequent block. Analyses revealed a significant group main effect ($F_{(3,124)} = 8.499$, $p < 0.001$, partial $\eta^2 = 0.171$), but no rest block ($F_{(9.716,1204.784)} = 1.280$, $p = 0.239$, partial $\eta^2 = 0.010$) nor a rest block x group interaction effect ($F_{(29.148,1204.784)} = 0.957$, $p = 0.532$, partial $\eta^2 = 0.023$; Figure S6B). Pairwise comparisons on the group main effect indicated that children exhibited significantly larger micro-offline gains as compared to adolescents ($F_{(1,62)} = 6.877$, $p = 0.011$, partial $\eta^2 = 0.100$), young ($F_{(1,61)} = 13.965$, $p < 0.001$, partial $\eta^2 = 0.186$) and older adults ($F_{(1,61)} = 11.436$, $p = 0.001$, partial $\eta^2 = 0.158$; Figure S6D).

In summary, the calculation of the micro-online and -offline gains based on the non-normalized data demonstrates that children exhibited smaller and larger micro-online and -offline performance gains, respectively compared to the other 3 age groups.

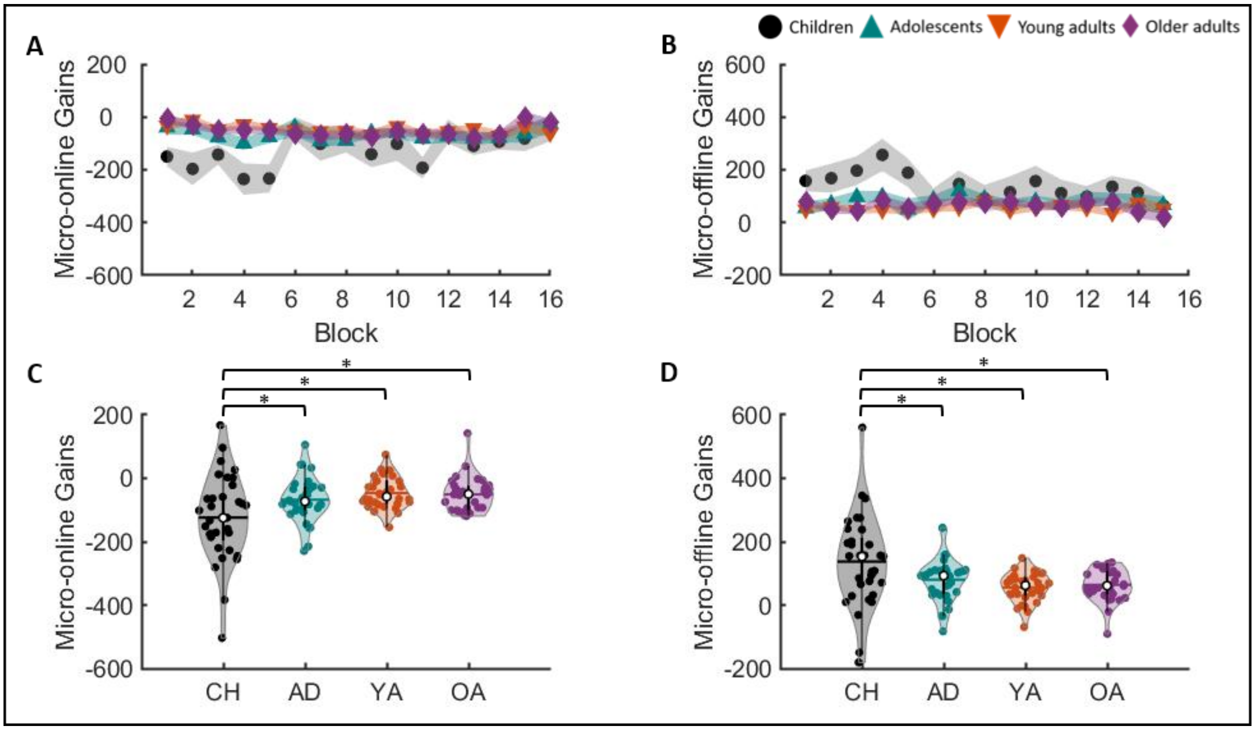

***Figure S6.*** *Micro-online (**left**) and -offline (**right**) gains. Panels A and B depict micro gains displayed as a function of practice blocks. Panels C and D contain violin plots of micro-online and -offline, respectively, gains averaged across blocks. Note that pairwise comparisons in these panels included blocks or rest periods in the statistical models. Shaded regions represent the kernel density estimate of the data, colored circles depict individual data, open circles represent group medians, and the horizontal lines depict group means[9].* $p < 0.05$ for differences between pairs of groups.*

### Appendix 4: Assessment of macro-offline performance changes with non-normalized data (Experiment 2)

Macro-offline performance changes in the main text were based on inter-session differences in *normalized* performance outcomes, adjusted for the baseline performance on the pre-learning random run. Here, we conduct exploratory analyses on macro-offline performance changes computed on non-normalized response times (RTs).

*4.1. Sequential macro-offline performance changes*

The sequential macro-offline gains were re-assessed with the same computation as described in the main text but using the non-normalized RTs (see Figure S7). Results revealed both offline period ($F_{(1,104)}$ = 94.16, p < 0.001, partial $\eta^2$ = 0.475) and group ($F_{(3,104)}$ = 18.02, p < 0.001, partial $\eta^2$ = 0.342) main effects, as well as a significant offline period x group interaction effect ($F_{(3,104)}$ = 5.62, p = 0.001, partial $\eta^2$ = 0.140). Follow-up comparisons revealed significant group effects for both the 5-hour ($F_{(3,107)}$ = 6.12, p < 0.001, $\eta^2$ = 0.150) and the 24-hour ($F_{(3,107)}$ = 24.71, p < 0.001, $\eta^2$ = 0.416) sequential macro-offline gains. Post-hoc pairwise comparisons indicated that the sequential macro-offline gains after a 5-hour period of wakefulness were higher in children and adolescents as compared to the young (p = 0.018, G = 0.632; p = 0.061, G = 0.563, respectively) and older adults (p = 0.004, G = 1.254; p = 0.018, G = 1.291, respectively). The children also showed higher 24-hour macro-offline gains in comparison to the adolescents (p = 0.001, G = 0.835), young (p < 0.001, G = 1.575) and older (p < 0.001, G = 2.185) adults. The older adults displayed significantly lower 24-hour macro-offline gains than all the other age groups (adolescents: p < 0.001, G = 1.183; young adults: p = 0.058, G = 1.015). The findings above are consistent with the results based on the normalized data presented in the main text.

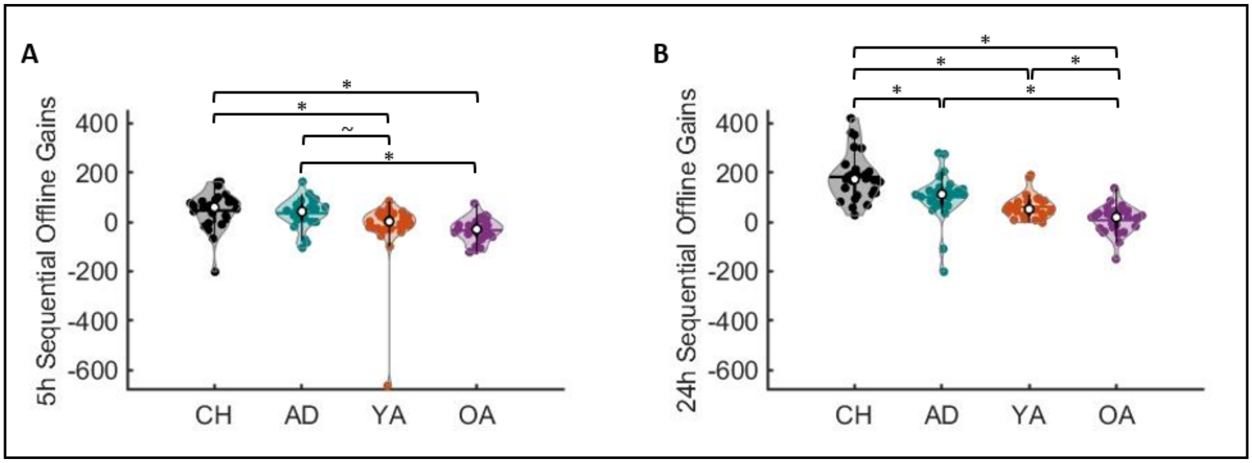

***Figure S7.*** *Non-normalized sequential offline gains across the 5-hour (**Panel A**) and 24-hour (**Panel C**) offline periods for the four age groups. Shaded regions represent the kernel density estimate of the data, colored circles depict individual data, open circles represent group medians, and the horizontal lines depict group means[9]. * p < 0.05 and ~ p < 0.10 for pairwise group comparisons.*

*4.2. Random and sequence-specific macro-offline performance gains*

Macro-offline gains in the random condition were assessed with the same computation as described in the main text but using the non-normalized RTs (Figure S8A). Significant main effects for both offline period ($F_{(1,103)} = 15.759$, $p < 0.001$, partial $\eta^2 = 0.133$) and group ($F_{(3,103)} = 3.474$, $p = 0.019$, partial $\eta^2 = 0.092$) were revealed, but there was no offline period x group interaction effect ($F_{(3,103)} = 1.135$, $p = 0.339$, partial $\eta^2 = 0.032$). Post-hoc pairwise comparisons on group differences across the two offline intervals indicated that older adults exhibited smaller random macro-offline gains as compared to children ($p = 0.017$, $G = 0.656$).

Sequence-specific macro-offline gains were assessed with the same computation as described in the main text but using with non-normalized RTs (Figure S8B). Significant main effects for both offline period ($F_{(1,103)} = 32.333$, $p < 0.001$, partial $\eta^2 = 0.239$) and group ($F_{(3,103)} = 7.523$, $p < 0.001$, partial $\eta^2 = 0.180$) were revealed, but there was no offline period x group interaction effect ($F_{(3,103)}$

= 2.079, p = 0.108, partial $\eta^2 = 0.057$). Post-hoc pairwise comparisons on group differences across the two offline intervals indicated that children exhibited larger sequence-specific macro-offline gains as compared to young (p = 0.026, G = 0.627) and older adults (p < 0.001, G = 0.949). Additionally, adolescents showed significantly larger sequence-specific offline gains in comparison to older adults (p = 0.003, G = 1.360). These results suggest that the offline changes in children likely involved the strengthening of their sequential memory. These findings are consistent with the results based on the non-normalized RTs presented in Appendix 8 below.

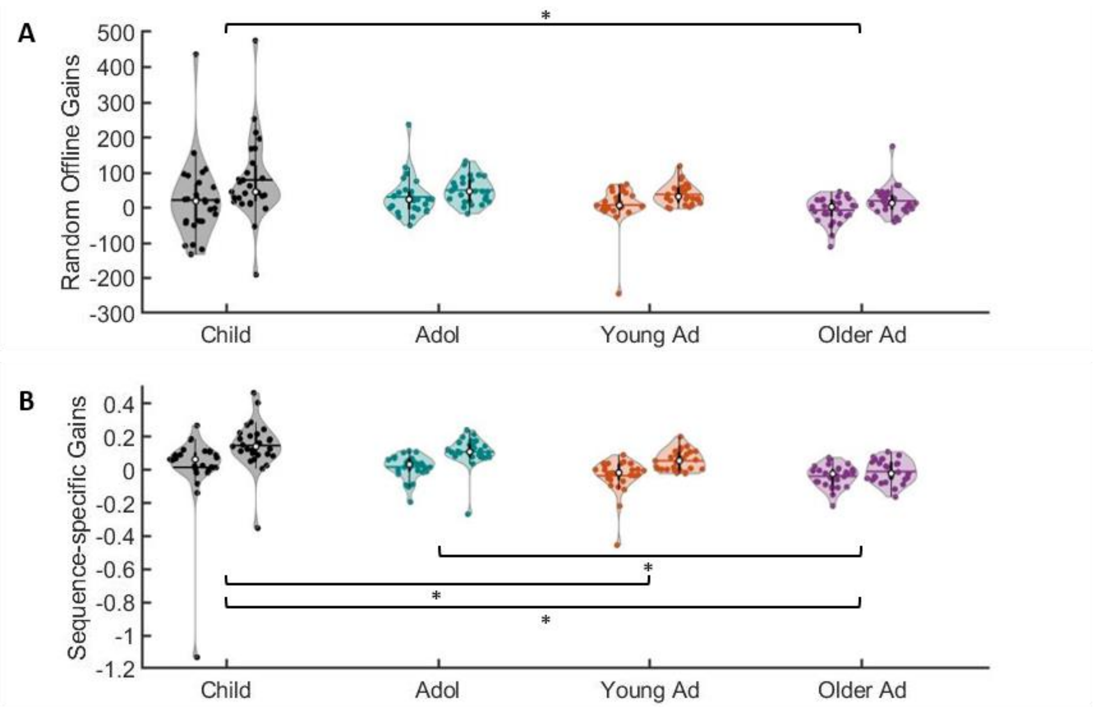

***Figure S8**: **A)** Non-normalized random offline gains across the 5-hour (left plots) and 24-hour (right plots) offline periods plotted per age group. **B)** Non-normalized sequence-specific offline gains across the 5-hour (left plots) and 24-hour (right plots) offline periods. Shaded regions represent the kernel density estimate of the data, colored circles depict individual data, open circles represent group medians, and the horizontal lines depict group means[9].*

### Appendix 5: Assessment of initial learning dynamics with normalized performance measures (Experiment 2).

***Table S5**: Results from statistical analyses on normalized task performance measures of session 1 in Experiment 2.*

| | RT | | | | Accuracy | | | |
|---|---|---|---|---|---|---|---|---|
| **Effect** | **df** | **F** | **p** | **Part $\eta^2$** | **df** | **F** | **p** | **Part $\eta^2$** |
| *A. Training* | | | | | | | | |
| Block | 7.87,817.92 | 48.34 | <0.001* | 0.317 | 6.27,651.7 | 0.861 | 0.527 | 0.008 |
| Group | 3,104 | 1.57 | 0.201 | 0.043 | 3,104 | 1.09 | 0.356 | 0.031 |
| B x G | 23.59, 817.92 | 1.10 | 0.341 | 0.031 | 18.80,651.7 | 1.03 | 0.419 | 0.029 |
| *B. Test* | | | | | | | | |
| Block | 2.68,278.3 | 0.25 | 0.837 | 0.002 | 2.25,233.48 | 2.25 | 0.101 | 0.021 |
| Group | 3,104 | 0.64 | 0.592 | 0.018 | 3,104 | 4.31 | 0.007* | 0.111 |
| B x G | 8.03,278.3 | 1.37 | 0.211 | 0.038 | 6.74,233.48 | 1.64 | 0.129 | 0.045 |
| *C. Post-learning random* | | | | | | | | |
| Block | 2.38,247.9 | 1.60 | 0.200 | 0.015 | 2.14,222.51 | 1.86 | 0.155 | 0.018 |
| Group | 3,104 | 2.30 | 0.082 | 0.062 | 3,104 | 0.76 | 0.522 | 0.021 |
| B x G | 7.15,247.9 | 1.10 | 0.365 | 0.031 | 6.42,222.51 | 0.83 | 0.558 | 0.023 |

*Results from statistical analyses of normalized response time (RT) and accuracy measures for all task runs of session 1 of Experiment 2. Significant values are marked with an asterisk. Df = degrees of freedom; part $\eta^2$ = partial eta squared; B x G = block x group interaction. Figure 4 in the main text depicts the learning dynamics of these normalized measures.*

### Appendix 6: Relationships between micro and macro-offline performance changes (Experiment 2)

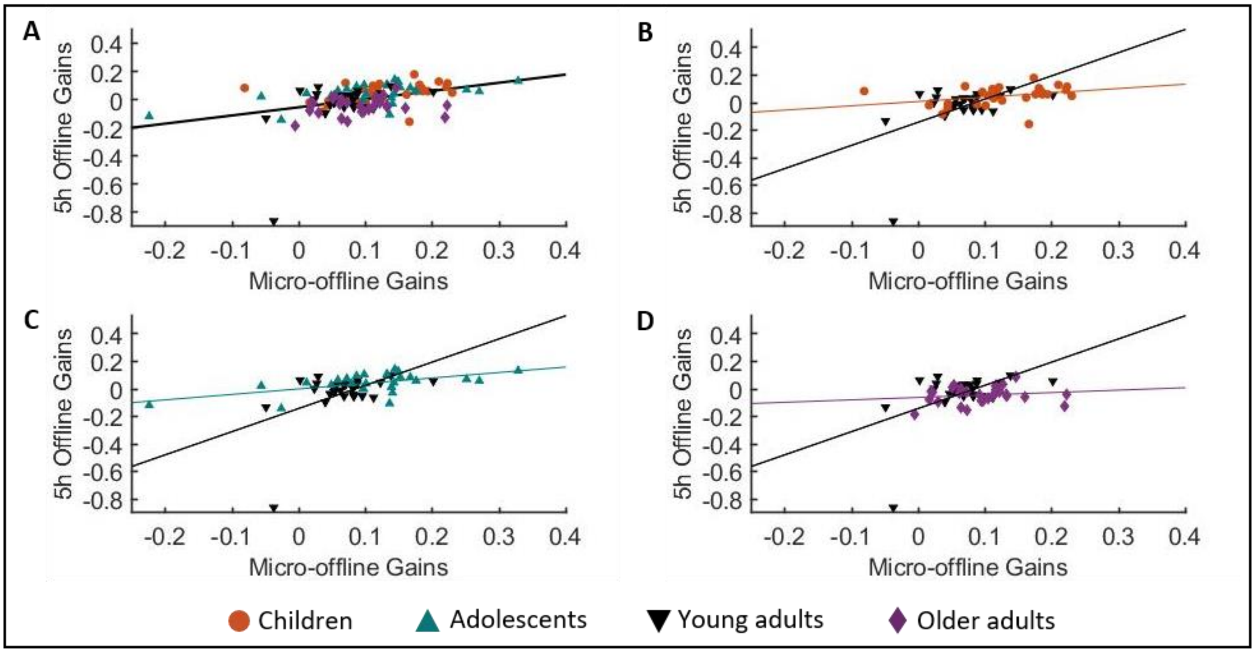

***Figure S9**: Five-hour macro-offline gains plotted as a function of the micro-offline gains (data points are color coded by group).* In ***panel A,*** *the regression line fits data collapsed across the 4 age groups. Panels B-D compare young adults to children (B), adolescents (C) and older adults (D). Details of the corresponding statistical results are presented in the main text. In brief, there was a significant relationship between micro- and macro-offline gains collapsed across the 4 age groups (Panel A). Between group comparisons revealed that this relationship was significantly different between young adults and each of the other 3 age groups. Note, however, that these between-group differences were no longer significant if the young adult with the large negative 5hr offline gains was excluded from analyses.*

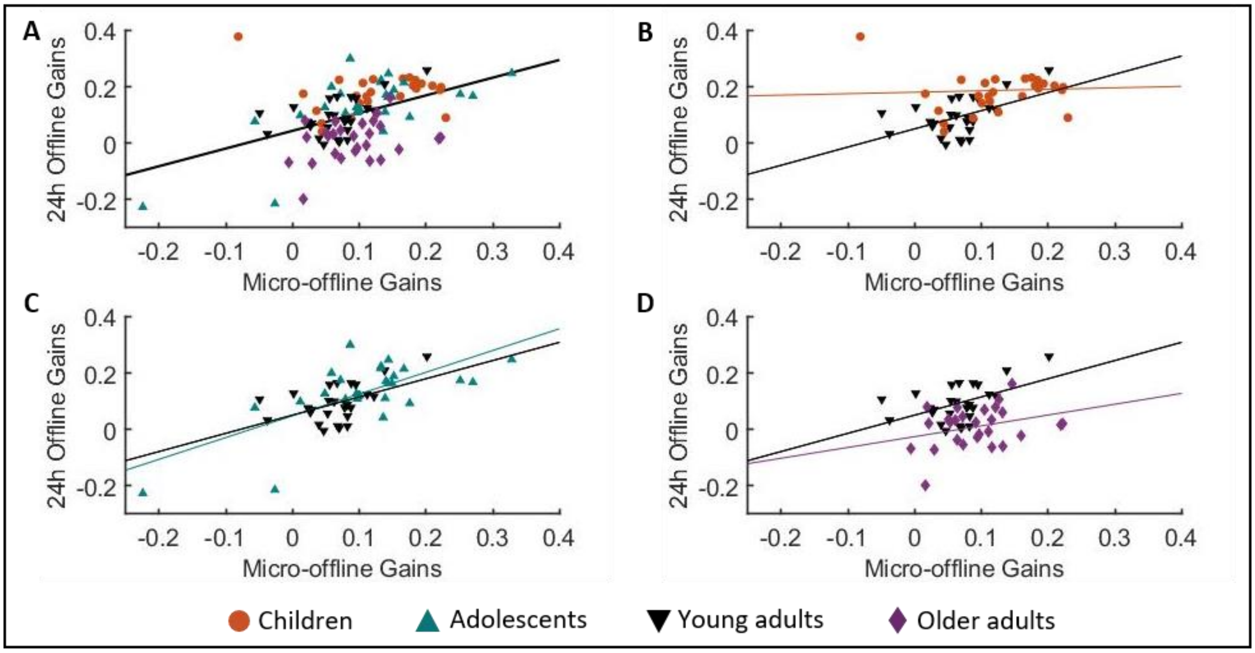

***Figure S10**: Twenty-four-hour macro-offline gains plotted as a function of the micro-offline gains (data points are color coded by group). In **panel A,** the regression line fits data collapsed across the 4 age groups. Panels B-D compare young adults to children (B), adolescents (C) and older adults (D). Corresponding statistical results are presented in the main text. In brief, there was a significant positive relationship between micro-offline and 24hr macro-offline gains collapsed across groups. This relationship was significantly different between young adults and children (Panel B) but not between young adults and adolescents (Panel C) or older adults (Panel D).*

### Appendix 7: One-sample t-tests assessing dependent measures of interest (Experiments 1 and 2)

***Table S6****: Results of one-sample t-tests for Experiment 1*

| Variable | df | t | p | G |
|---|---|---|---|---|
| *A. Learning magnitude* | | | | |
| Children | 32 | 4.04 | < 0.001* | 0.686 |
| Adolescents | 32 | 6.87 | < 0.001* | 1.168 |
| Young adults | 31 | 8.80 | < 0.001* | 1.518 |
| Older adults | 31 | 7.39 | < 0.001* | 1.275 |
| *B. Micro-online gains* | | | | |
| Children | 32 | -5.82 | < 0.001* | -0.990 |
| Adolescents | 32 | -5.58 | < 0.001* | -0.948 |
| Young adults | 31 | -5.47 | < 0.001* | -0.943 |
| Older adults | 31 | -6.01 | <0.001* | -1.037 |
| *C. Micro-offline gains* | | | | |
| Children | 32 | 6.14 | < 0.001* | 1.043 |
| Adolescents | 32 | 6.75 | < 0.001* | 1.147 |
| Young adults | 31 | 6.89 | < 0.001* | 1.188 |
| Older adults | 31 | 8.01 | < 0.001* | 1.381 |

*Results of one-sample t-tests to determine and statistically test directionality of several variables of experiment 1. Test value = 0. Significant values are marked with an asterisk; Df = degrees of freedom; G = Hedge's correction; The interpretation of relevant comparisons can be found in the discussion of the main text.*

***Table S7**: Results of one-sample t-tests for Experiment 2*

| **Variable** | **df** | **t** | **p** | **G** |
|---|---|---|---|---|
| *A. 5-hour sequential offline gains* | | | | |
| Children | 26 | 3.49 | 0.002* | 0.653 |
| Adolescents | 26 | 3.05 | 0.005* | 0.570 |
| Young adults | 26 | -0.94 | 0.356 | -0.175 |
| Older adults | 26 | -3.85 | < 0.001* | -0.719 |
| *B. 24-hour sequential offline gains* | | | | |
| Children | 26 | 13.84 | < 0.001* | 2.586 |
| Adolescents | 26 | 5.83 | < 0.001* | 1.089 |
| Young adults | 26 | 7.48 | < 0.001* | 1.398 |
| Older adults | 26 | 0.63 | 0.538 | 0.117 |

*Results of one-sample t-tests to determine and statistically test directionality of the sequential macro-offline gains of experiment 2. Test value = 0. Significant values are marked with an asterisk; Df = degrees of freedom; G = Hedge's correction; The interpretation of relevant comparisons can be found in the discussion of the main text.*

### Appendix 8: Random and sequence-specific macro-offline performance changes with normalized performance measures (Experiment 2)

Figure S11A displays the random macro-offline gains across both offline intervals per age group and as a function of age. Significant main effects for both offline period ($F_{(1,103)} = 21.409$, $p < 0.001$, partial $\eta^2 = 0.172$) and group ($F_{(3,103)} = 4.097$, $p = 0.009$, partial $\eta^2 = 0.107$) were revealed, but there was no offline period x group interaction effect ($F_{(3,103)} = 0.601$, $p = 0.616$, partial $\eta^2 = 0.017$). Post-hoc pairwise comparisons on group differences across the two offline intervals indicated that older adults exhibit lower random macro-offline gains as compared to children ($p = 0.023$, $G = 0.700$) and adolescents ($p = 0.012$, $G = 1.041$). Importantly, the children did not differ from any of the other age groups.

Figure S11B displays the sequence-specific macro-offline gains across both offline intervals per age group and as a function of age. Significant main effects for both offline period ($F_{(1,103)} = 32.558$, $p < 0.001$, partial $\eta^2 = 0.240$) and group ($F_{(3,103)} = 7.210$, $p < 0.001$, partial $\eta^2 = 0.174$) were revealed, but there was no offline period x group interaction effect ($F_{(3,103)} = 1.737$, $p = 0.164$, partial $\eta^2 = 0.048$). Post-hoc pairwise comparisons on group differences across the two offline intervals indicated that children exhibited larger sequence-specific macro-offline gains as compared to young ($p = 0.034$, $G = 0.609$) and older adults ($p < 0.001$, $G = 0.923$). Additionally, adolescents showed significantly larger sequence-specific offline gains in comparison to older adults ($p = 0.003$, $G = 1.340$).

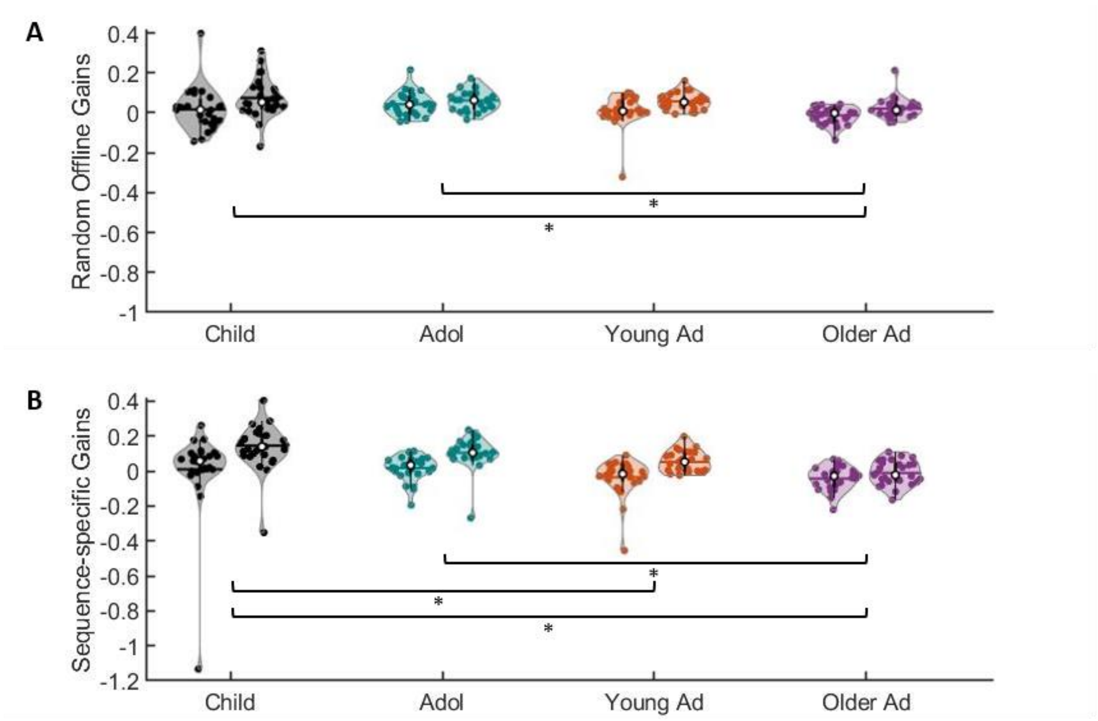

***Figure S11**: **A)** Random offline gains across the 5-hour (left plots) and 24-hour (right plots) offline periods plotted per age group. **B)** Sequence-specific offline gains across the 5-hour (left plots) and 24-hour (right plots) offline periods. Shaded regions represent the kernel density estimate of the data, colored circles depict individual data, open circles represent group medians, and the horizontal lines depict group means*[9]
